# TIF1γ drives oral cancer recurrence by the transcriptional regulation of self-renewal genes, such as *HES1*

**DOI:** 10.64898/2025.12.02.691793

**Authors:** K P Padmaja, Amrutha Mohan, G Madhumathy Nair, Jyothi S Prabhu, Hafsa Shabeer, V P Snijesh, P G Balagopal, A Thameem, Alan Jose, Nebu Abraham George, Jiss Maria Louis, Parvathy Visakh, Gayathri V Nair, Anna P Joseph, Vishnu Sunil Jaikumar, Riya Ann Paul, Jackson James, Tessy Thomas Maliekal

**Author notes:** Equal contributing authors (K P Padmaja, Amrutha Mohan).

## Abstract

TIF1γ is an E3 ubiquitin ligase and key mediator of the noncanonical TGF-β signaling pathway. Initially characterized for its developmental functions, TIF1γ is essential for maintaining the pluripotency of adult stem cells, including long-term hematopoietic stem cells. Although TGF-β signaling contributes to cancer progression and recurrence, TIF1γ has traditionally been regarded as a tumor suppressor due to its inhibition of oncogenes involved in epithelial–mesenchymal transition. However, clinical reports have associated high TIF1γ expression at advanced cancer stages with poor prognosis. To elucidate the mechanism underlying this paradox, we identify a previously unrecognized role of TIF1γ in promoting the self-renewal capacity of oral cancer cells, thereby contributing to disease recurrence. Phosphoproteomic profiling of self-renewal–enriched cells revealed activation of a noncanonical TGF-β pathway. Using extreme limiting dilution assays, an ALDH1A1-DsRed2 cancer stem cell reporter, multiple oral cancer cell lines, primary 3D cultures on alginate matrix, and orthotopic mouse models, we demonstrate that TIF1γ depletion significantly reduces self-renewal and prolongs disease-free survival. Immunoprecipitation (IP)–LC/MS/MS analysis identified transcriptional regulators within the TIF1γ interactome, including TRRAP and H2A.Z, which were validated by IP and FRET assays. ChIP and IP studies further revealed that during self-renewal enrichment, TRRAP acetylates H2A.Z, decreasing the chromatin occupancy of its unacetylated form. Acetylated H2A.Z is subsequently recognized by TIF1γ, which monoubiquitinates H2B at promoters of self-renewal genes such as *HES1*, initiating transcription. In alignment with findings from mouse neocortical development, where a Notch-independent *Hes1*-expressing (NIHes1) population defines primitive quiescent stem cells, we show that TIF1γ acts as an acetylation reader specifically at the NIHES1 promoter region of *HES1*. TIF1γ depletion drives NIHES1 cells toward a Notch-dependent *HES1* (NDHES1) identity. RNA-seq confirmed reversal of 100 NIHES1-specific genes following TIF1γ loss, along with downregulation of pluripotency-associated genes found in embryonic stem cells, supporting a critical role for TIF1γ in maintaining primitive cancer stem cell states. Consistent with our *in vivo* findings, primary oral cancer samples showed that increased frequencies of TIF1γ^+^/TRRAP^+^/H2A.Z^-^ cells strongly predict recurrence. Given that the histone acetylation–reader function of TIF1γ drives poor prognosis, our findings suggest the TIF1γ bromodomain as a potential therapeutic target requiring further investigation.

## INTRODUCTION

Transforming growth factor-β (TGF-β), a pleotropic cytokine that regulates cell growth, differentiation, apoptosis, and homeostasis, can fuel disease progression, when deregulated. TGF-β exerts a dual effect in cancer- while it acts as a tumor suppressor in the initial stages, it promotes cancer progression and metastasis in the later stages of the disease (1, 2). The ligand-dependent activation of Smad2/3 by the TGF-β receptor I (ALK5) initiates the canonical TGF-β pathway leading to the transcriptional regulation of target genes by the active Smad2/3/4 complex. At the same time, many TGF-β target genes are activated independent of Smads, when TGF-β receptors interact with intermediates of other signaling pathway, initiating a non-canonical signaling (2, 3). Transcriptional Intermediary Factor 1γ or TIF1γ was identified as an intermediary molecule of non-canonical TGF-β signaling, involved in hematopoiesis (4). Later, TIF1γ was shown to regulate Wnt (5), Androgen receptor (6) and Estrogen receptor signaling (7).

TIF1γ, also known as TRIM33, RFG7, PTC7 or Ectodermin, is a known E3 Ubiquitin ligase. This molecule was initially reported as an E3 Ubiquitin ligase for Smad4, a signal transducer of TGF-β signaling, thereby inducing ectoderm specification in Xenopus embryos (8). Subsequently, its E3 Ubiquitin ligase activity was well explored in the context of development and cancer. This E3 Ubiquitin ligase activity plays a role in embryonic development (8), erythroid differentiation (9, 10), DNA repair (11) and mitosis (12). In addition to its E3 Ubiquitin ligase activity, TIF1γ is a transcriptional regulator acting as a transcriptional activator or repressor in different contexts. One of the first reports showing TIF1γ as a transcriptional repressor was in adult hematopoiesis, where it regulates the process through the E3- ubiquitin ligase-independent repression of TAL1 and PU.1. target genes (13). Further, it acts as a transcriptional repressor in dendritic cell differentiation (14) and in the exit of totipotency of two-cell stage embryo (15). On the other hand, in mouse embryonic stem cells, TIF1γ co-localizes with PML on the active promoters of nodal target genes such as Lefty1/2, suggesting it as a transcriptional activator (16). The E3-independent transcriptional activator role of TIF1γ is also ascertained in Th17 differentiation (17). Moreover, estrogen receptor (ER) and androgen receptor (AR) recruit TIF1γ to the promoters of their target genes, establishing their transcriptional activator role (6, 7). Collectively, the existing evidences reinforces the transcriptional regulatory role of TIF1γ, though the exact mechanism is not well-characterized.

The quest to find the role of TIF1γ in cancer began with its identity as a tumor suppressor in pancreatic cancer. The study showed that the loss of TIF1γ cooperated with Kras mutations in the pathogenesis of cystic pancreatic tumors. Consistent with that, TIF1γ expression is down-regulated in pancreatic cancer samples, compared to normal pancreatic tissues (18, 19). The expression of TIF1γ is also down regulated in squamous cell carcinoma, lung adenocarcinoma, endometrial carcinoma, clear cell renal cell carcinoma and hepatocellular carcinoma, suggesting its tumor suppressor role. Mechanistically, this tumor suppression is attributed to the E3 Ubiquitin ligase-dependent suppression of oncogenic molecules such as Smad4, SnoN1, Myc, and β-Catenin (20–23). Paradoxically, TIF1γ is shown to associate with poor prognosis in breast cancer and colorectal cancer, suggesting a protumorigenic function of TIF1γ (24–26). Later, its tumor promoter function was reported in estrogen receptor (ER)α-positive breast cancer (7) and prostate cancer (6). In prostate cancer, TIF1γ, by virtue of the RING domain E3 ubiquitin ligase activity, stabilizes androgen receptor by preventing another E3 ubiquitin ligase Skp2. Similarly, when TIF1γ is recruited to the promoter region of ER target genes, despite its E3 ubiquitin ligase activity, it stabilizes the receptor, leading to increased transcriptional output and estrogen–dependent cancer progress, though the mode of stabilization of ERα is unclear (7).

Since TIF1γ was identified as a regulator of hematopoiesis, its functional aspects are best studied in development than cancer. In mice, TIF1γ is required for the exit from totipotency (2C) stage to the pluripotency stage (15). Also, in the pluripotent embryoid bodies, TIF1γ is essential for further differentiation to mesoderm (27). Later, it was shown that TIF1γ is recruited to H3K18 acetylation marks of the promoters of mesendodermal genes for its transcription (28). So, in the early embryonal development, TIF1γ induces differentiation from totipotent cell to pluripotent embryoid bodies to mesendoderm to mesoderm. But in the case of adult pluripotent stem cells, such as hematopoietic stem cells (HSCs), the scenario is slightly different. Although TIF1γ deficient long term HSCs (LT-HSCs) were able to sustain hematopoiesis, its loss resulted in increased differentiation of LT-HSCs to short-term HSCs (ST-HSCs) and then to myeloid precursors, suggesting that TIF1γ is essential to maintain pluripotency of LT-HSCs (13). Though TIF1γ plays an inevitable role in the self-renewal of normal stem cells, its role in the regulation of self-renewal of cancer cells remains unexplored.

Akin to normal stem cells, cancer cells also exhibit self-renewal capability, which are known as cancer stem cells (CSCs) that are the primary drivers of unlimited proliferation of cancer cells. Self-renewal ability of cancer cells is the underlying cause of recurrence of the disease, which is often correlated to high mortality rate (29–31). Oral cancer, one of the leading causes of cancer deaths worldwide is a major health problem in Indian subcontinent. According to GLOBOCAN 2022, oral cancer is the third most common cancer, in terms of incidence and mortality, in India. It is estimated that in India, one out of two oral cancer patients die within 5years after treatment (32, 33). The high mortality rate of oral cancer is significantly associated with the recurrence of the disease (32). As reported for many other cancers, the self-renewing CSC population is associated with recurrence and poor prognosis of oral cancer (34).

In the current study we elucidated the role of TIF1γ in recurrence or oral cancer, with special reference to its molecular mechanism regulating self-renewal of oral cancer cells. Since our preliminary data identified the enrichment of non-canonical TGF-β signaling in self-renewing conditions, we explored the possibility of TIF1γ, a mediator of non-canonical TGF-β signaling, as a regulator of self-renewal in oral cancer cells. Further, we investigated its transcriptional regulatory mechanism underlying this protumorigenic role. Our results revealed that the transcriptional regulatory function of TIF1γ is dependent both on its E3 Ubiquitin ligase activity and its histone acetylation reader activity.

## RESULTS

### Non-canonical TGF-β signaling supports self-renewal of oral cancer cells

Since self-renewing population is known to be the cause of recurrence, we performed a phosphoproteomic analysis to identify the signaling pathways supporting the enrichment of self-renewing population in oral cancer. The phosphoproteomic analysis of HSC-4 oral cancer cells revealed that TGF-β signaling pathway is activated in self-renewing population-enriched sphere condition, compared to monolayer condition. The molecules differentially phosphorylated are shown in Figure S1. To test whether TGF-β signaling is important for self-renewal, we used primary cell culture form Oral Squamous Cell Carcinoma (OSCC) patients. The CD44^+^ oral cancer cells were FACS sorted, and their keratinocyte origin was confirmed by pancytokeratin and EpCAM positivity (Figure S2A-D). Though the cancer cells express TGF-β, the major source for the same are stromal cells. So, in our experiment, a mat of mitomycin-arrested DPC/fibroblasts were taken as the source of TGF-β. We depleted TGF-β, either by a neutralizing antibody or by lentiviral knockdown (Figure S2E). When the primary OSCC cells were grown over the wild type DPC cells, the cells expressed the universal stem cell marker, ALDH1A1. At the same time, when TGF-β was depleted, the ALDH1A1 population decreased significantly in all the four samples (Figure 1A).

**Figure 1:**
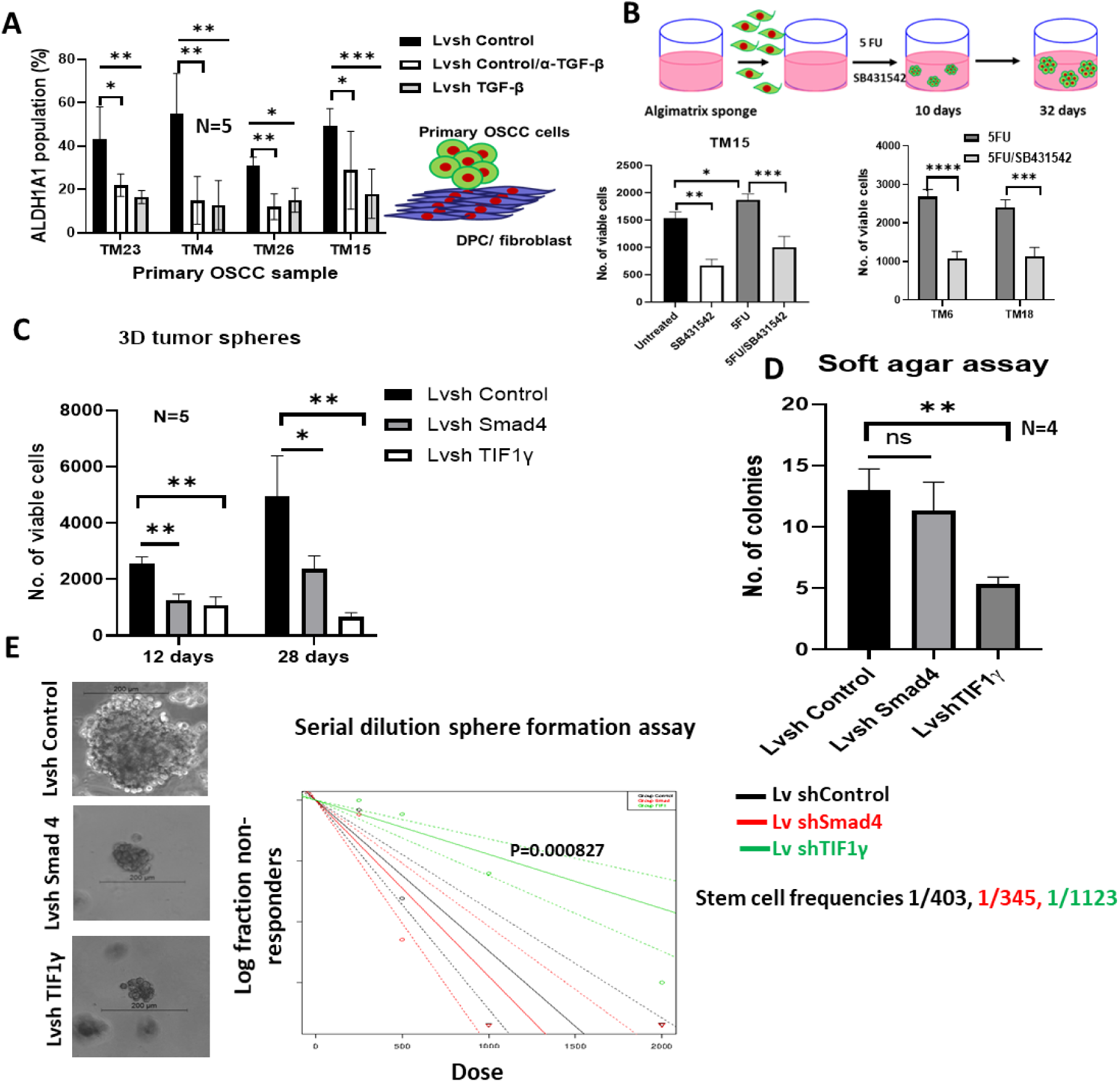
Non-canonical TGF-B signallingl?’gsgilates self-renewal. (A)OSCC sample keratinocytes were co-cultured on wild type DPC cells (control) or cells deficient of TGF-p (DPC-Lv-ShTGF-p). Keratinocytes grown on wild type were treated with neutralizing antibody for TGF-p (0.5 ug/ml). The cells were fixed with 4% PFA and immunostained using CD44-FITC and ALDHI1A1/Alexa 680. The percentage expression of ALDHIA1 was plotted after counting 10 fields in 60X with positive expression for ALDHIAL. Error bar represents the SD. (B,C) The CD44+ tumor cells and mitomycin arrested CD44- fibroblasts from the respective OSCC samples were used for 3D co-culture. The drug, 5-FU (10pg/ml) and SB431542 treatment (5puM) were started on 8 day and continued for 10 days. The surviving cells were allowed to grow for 10days and trypan blue exclusion assay was performed to evaluate the total viable cells. Error bar represents the SEM. (D) The control cells, Smadd-depleted cells and TIFly-depleted cells were grown for 28-Days as 3D culture and the matrix was dissolved at the indicated time point to count the number of viable cells after trypan blue exclusion. Error bar represents the SEML.(E) The control cells, Smad4-depleted cells and TIF1y-depleted cells were used for serial dilution sphere formation assay. The stem cell frequency was calculated using ELDA. The log fraction plot for the same is given.

Next, using an *in vitro* 3D culture model on alginate matrix using primary OSCC samples, we checked whether TGF-β is critical for recurrence after treatment with a chemotherapeutic drug used in oral cancer, 5-FU (Figure 1B). Though 5-FU induced cell death, after the withdrawal of the drug, surviving cells were able to proliferate more, compared to the untreated cells. But the inhibition of type I TGF-β receptor by the treatment of SB431542 drastically reduced the capability of 5-FU surviving cells to proliferate in all the three primary OSCC samples (Figure 1B).

When we analyzed the phosphoproteomic results, the canonical Smad mediators were not differentially phosphorylated in sphere condition (Figure S1), suggesting it to be a non-canonical TGF-β signaling. So, we next analyzed whether the canonical or non-canonical TGF-β signaling is involved in the induction of self-renewal. We chose Smad 4 as the canonical mediator and TIF1γ as the non-canonical mediator. 3D tumor spheres were cultured using Smad4 knocked-down, TIF1γ knocked-down and control cells (Figure S3A) and the cell viability was checked on 12^th^ and 28^th^ day. Our results showed that both Smad4 and TIF1γ are essential in keeping the tumor burden as there was a significant reduction in the viable cell count in the cells depleted of either of the molecules compared to control on both 12^th^ and 28^th^ day (Figure 1C). Of note, we observed a steady increase in tumor bulk in control and Smad4 knocked-down condition with time, while there was a decrease observed in TIF1γ knocked-down condition. As the absence of sustained proliferation with time indicates lack of self-renewal ability, we evaluated the same using self-renewal assays like soft agar assay and extreme limiting dilution assay (ELDA). The number of soft agar colonies formed in the control and Smad 4-knocked-down cells were comparable while, there was a drastic reduction in the number of colonies in TIF1γ-knocked-down cells (Figure 1D). Next, we performed a more robust self-renewal assay, ELDA with serial dilution sphere formation. Though the spheres formed in both the knockdown conditions were smaller compared to the control cells, the cancer stem cell (CSC) frequency was not significantly different in Smad 4-depleted cells compared to control cells (Figure 1E, Figure S3B). At the same time, the CSC frequency decreased by 2.7-fold upon TIF1γ knockdown (Figure 1E, Figure S3B). These results show that non-canonical TGF-β signaling through TIF1γ, and not the canonical pathway through Smad 4, plays a critical role in maintaining the self-renewing population in oral cancer.

### TIF1γ knockdown depletes self-renewing population in oral cancer

To explore the role of TIF1γ in self-renewal, we depleted TIF1γ in 4 different oral cancer cell lines, HSC-3, HSC-4, RCB1015, and RCB1017, using Lentiviral shRNA particles (Figure 2A, Figure S4A). *In vitro* serial dilution sphere formation assay showed that the stem cell frequency decreased by 2.6-fold in HSC-4, 1.9-fold in RCB1015 and 2.2-fold RCB1017 cell line upon TIF1γ knockdown when compared to the control (Figure 2B & S4C). Since pluripotency markers are enriched in CSCs, we analyzed the expression of pluripotent markers in these cells. There was a significant reduction in the mRNA levels of *OCT4, NANOG* and *KLF4* in TIF1γ-depleted RCB1015 and HSC-3 cell lines (Figure 2C, S5A & S5B). These *in vitro* results were further validated with an *in vivo* serial dilution xenograft assay using HSC-3 and RCB1015 cell lines.

**Figure 2:**
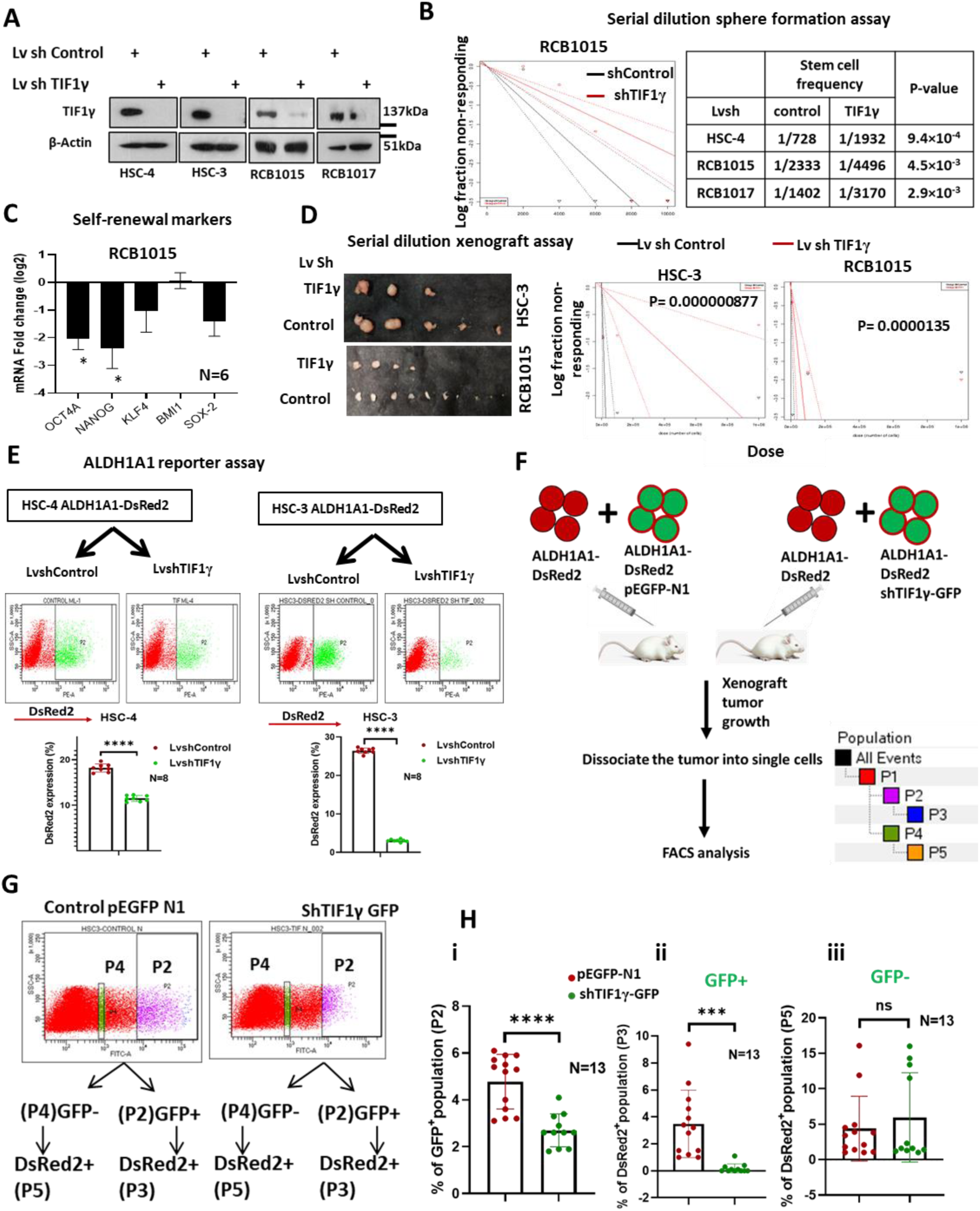
**TIF1γ , a non-canonical mediator of TGF-β signalling regulates self-renewal**. (A)Western blot experiment to show bands of TIF1γ in HSC-4, HSC-3, RCB1015 and RCB1017 oral cancer cells with knockdown of TIF1γ . β-Actin was used as the loading control. (B) The HSC-4,RCB1015 and RCB1017 cells with the TIF1γ knockdown and the control cells were used for serial dilution sphere formation assay. The stem cell frequency was calculated using ELDA. The representative graph of log fraction plot for RCB1015 is shown. (C) RCB1015 cells with the TIF1γ knockdown and the control cells grown as 3D spheres for 6 days were used for RNA isolation followed with qRT-PCR to analyze the expression of self-renewal genes. Log2 fold change normalized with *β-ACTIN* was plotted (D) The RCB1015 and HSC-3 TIF1γ knocked-down cells and control cells were serially diluted to 10 ,10 and 10 numbers and xenografts were generated for 21 days. The image for the tumors collected in 10 dilution or 10 dilutions of HSC-3 and RCB1015, respectively are shown. Log fraction plot of ELDA analysis is shown.(E) The FACS profile shows the percentage of ALDH1A1-DsRed2 population in TIF1γ knockdown and the control cells grown as monolayer. The error bars indicate standard deviation. (F) The HSC-4 ALDH1A1-DsRed2 cells and HSC-4 ALDH1A1-DsRed2 cells with TIF1γ knockdown tagged with GFP were mixed in 1:1 ratio and injected subcutaneously on the flanks of mice to generate a xenograft model. The wildtype and HSC-3 ALDH1A1-DsRed2 pEGFP-N1 was taken as a control. The tumor was collected and dissociated into single cells by enzymatic digestion. GFP+(P2) cells and GFP-(P4) cells were analysed for ALDH1A1- DsRed2 positivity (P3 and P5 population respectively). Population hierarchy is given. (G) Representative FACS analysis profile of GFP+ population in control and TIF1γ knockdown xenograft tumor (H) Quantitative representation of P2(i), P3(ii) and P5(iii). Error bar represents the SD.

The CSC frequency decreased significantly in TIF1γ-depleted HSC-3 and RCB1015 cell line-derived xenografts (Figure 2D, Figure S6A). To support these functional assays, we did some quantitative analyses of the CSC pool using reporter cell lines of CSCs harboring ALDH1A1-DsRed2 (35). TIF1γ was knocked-down in HSC-3 and HSC-4 cells harboring ALDH1A1-DsRed2 reporter construct (Figure S7A). A FACS analysis was done to quantify ALDH1A1+ve CSCs. There was a significant decrease in the ALDH1A1-DsRed2 expressing population in TIF1γ knocked-down cells compared to control cells in both cell lines (Figure 2E), indicating the role of TIF1γ in maintaining CSC pool. Taken together, our results show that cells losing TIF1γ expression loses the self-renewal ability.

As we expect heterogeneity in the expression level of TIF1γ among cancer cells, we checked whether TIF1γ imparts any survival advantage to cancer cells in a given tumor microenvironment. To recreate the heterogeneity with respect to TIF1γ expression, we mixed the wild type and TIF1γ-depleted HSC-3 ALDH1A1-DsRed2 cells in 1:1 ratio, and used them for xenograft development in immunocompromised mice (Figure 2F). Since the shRNA used for knockdown of TIF1γ had a GFP tag, we used empty vector with GFP tag as the control. The knockdown was confirmed by western blotting (Figure S7B). The xenograft tumors were collected and the cell suspension were analyzed by FACS for the GFP population (Figure 2G-H). The GFP population was significantly diminished in TIF1γ knocked-down cells compared to the control, indicating the inability of TIF1γ-depleted cells to survive in a tumor microenvironment (Figure2G & 2H). To confirm whether the depletion of TIF1γ deficient cells is due to the lack of self-renewal ability, we analyzed the DsRed2 population in the GFP+ (TIF1γ knockdown/wild type) and GFP- cells from both groups (both representing wild type). The cells lacking TIF1γ had significantly lesser DsRed2 expressing population among the GFP+ cells, while the wild type GFP- cells showed comparable DsRed2+ population (Figure2G & 2H). Taken together, our results showed that loss of TIF1γ impairs the self-renewal ability of cancer cells, and those TIF1γ-deficient cells will gradually diminish in a tumor microenvironment.

### TIF1γ depletion reduces chance of recurrence in mouse model

Since TIF1γ is regulating self-renewal, the expression of TIF1γ might eventually lead to the recurrence of the disease. In order to validate its role in recurrence, an *in vivo* orthotopic model of oral cancer was used (Figure 3A). The tumors were surgically removed, and the complete resection was ensured by bioimaging. The excised tumors from both the groups were of similar size and weight. However, the luciferase activity reflecting the viable tumor cells was significantly lower in the TIF1γ-depleted group (Figure 3B, C and D and Figure S8A). To understand why this discrepancy between luciferase flux and tumor size, we did a Hematoxylin & Eosin staining on the tumor sections. As the representative images shown (Figure 3E), TIF1γ-depleted tumors were mostly necrotic, while the control tumors had live tumor cells. This explained the disagreement between tumor size and luciferase flux, as only the live cells show luciferase activity. The animals were imaged at regular intervals post-surgery to check recurrence. On 8^th^ day post-surgery 6 animals from the control group and only one animal from the TIF1γ knocked-down group showed presence of tumor by *in vivo* imaging (Figure 3F, Figure S8A to E). While all the animals in the control group showed recurrence by 33^rd^ day, a half of the animals in the TIF1γ-depleted group did not show relapse till the 40^th^ day (Figure S8D&E). A Kaplan-Meier survival analysis revealed that the disease-free survival of the animals injected with TIF1γ-depleted cells is significantly higher compared to the control group, confirming the role of TIF1γ in recurrence (Figure 3G). Moreover, the disease progression in animals in the TIF1γ-deficinet group was significantly slower than that of the control group (Figure 3H). A reduced expression of ALDH1A1 and OCT4A observed on the xenograft sections of TIF1γ-depleted group compared to control confirmed that the better disease-free survival observed in the TIF1γ knocked-down group is indeed due to the loss of self-renewing population (Figure S9A). In summary, our mouse model analysis confirmed the role of TIF1γ in the regulation of self-renewal of oral cancer cells and its positive correlation to worse prognosis.

**Figure 3:**
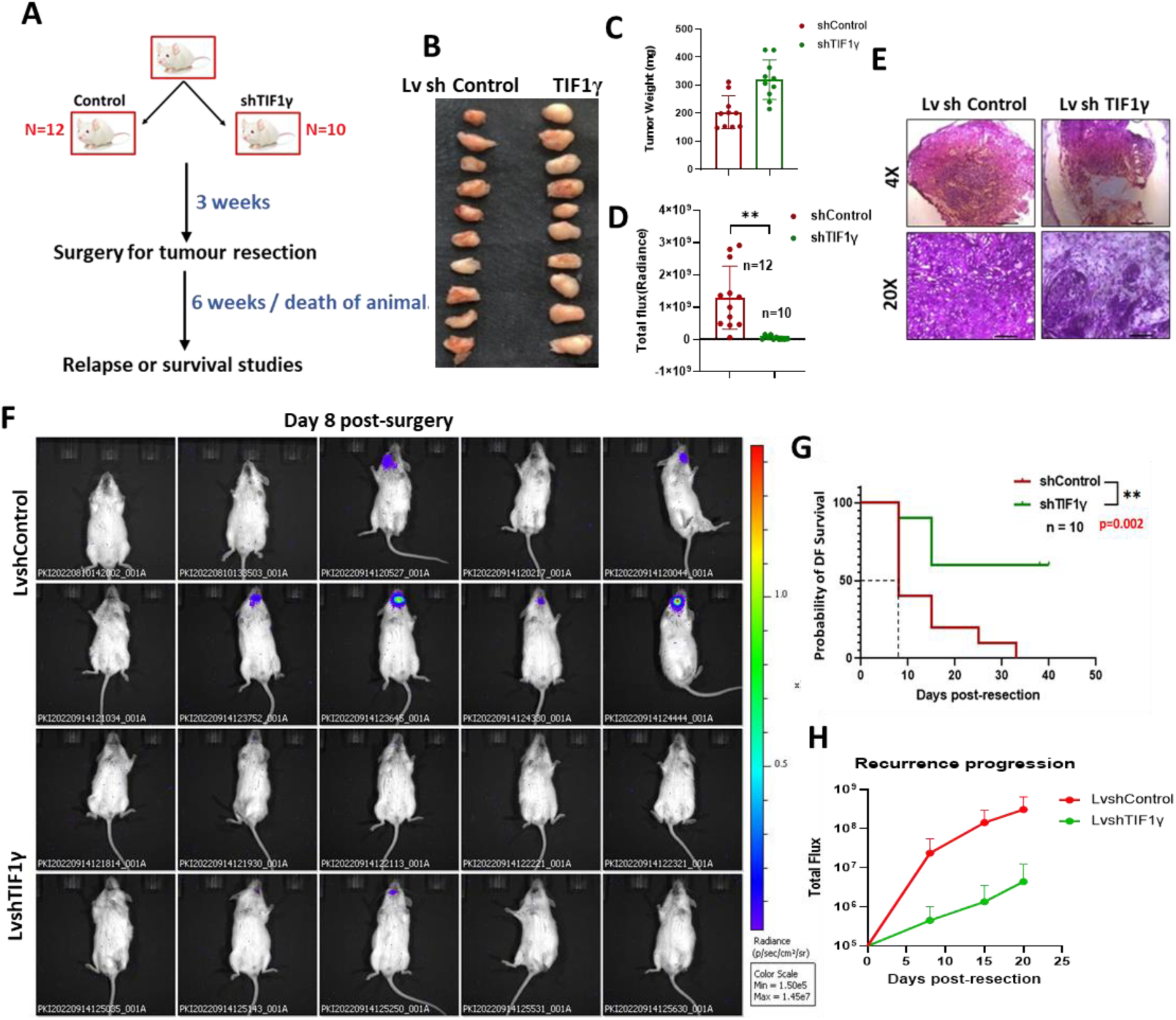
TIF1y regulates recurrence. (A) Schematic representation of the recurrence model. HSC-4 cells with stable luciferase expression with or without TIF1y knockdown were injected orthotopically into the NSG mice. The growth of the tumour was tracked by bio-imaging. (B) The image shows the tumours collected during the resection surgery on 21% day. (C) The histogram shows the tumour weight. Error bar represents the SD. (D)The total luciferase flux (measured in radiance) before the resection surgery. Error bar represents the SD. (E)The representative image for H&E staining for the tumours collected. The scale bar represents 500um and 100pum in 4X and 20X magnification respectively. (F) The luciferase activity were monitored at regular intervals post surgery to check relapse. (G) Detectable flux was used as end point and a Kaplan-Meier survival plot showing disease free survival of the animals post surgery with time was plotted. (H)The luciferase flux of animals till 20 day post-surgery is plotted. Error bar represents the SD.

### TIF1γ interacts with TRRAP and H2A.Z

Although all our experiments reinforced the notion that TIF1γ is a critical regulator of self-renewal ability and the chance of recurrence in oral cancer, its molecular mechanism remains uncertain. TIF1γ is a known E3 Ubiquitin ligase, which targets its substrates for proteasomal degradation. To identify the interacting partners of TIF1γ, an immunoprecipitation (IP) in presence of the proteasomal inhibitor, MG-132 was done followed by an LC/MS/MS analysis (Figure 4A).

**Figure 4:**
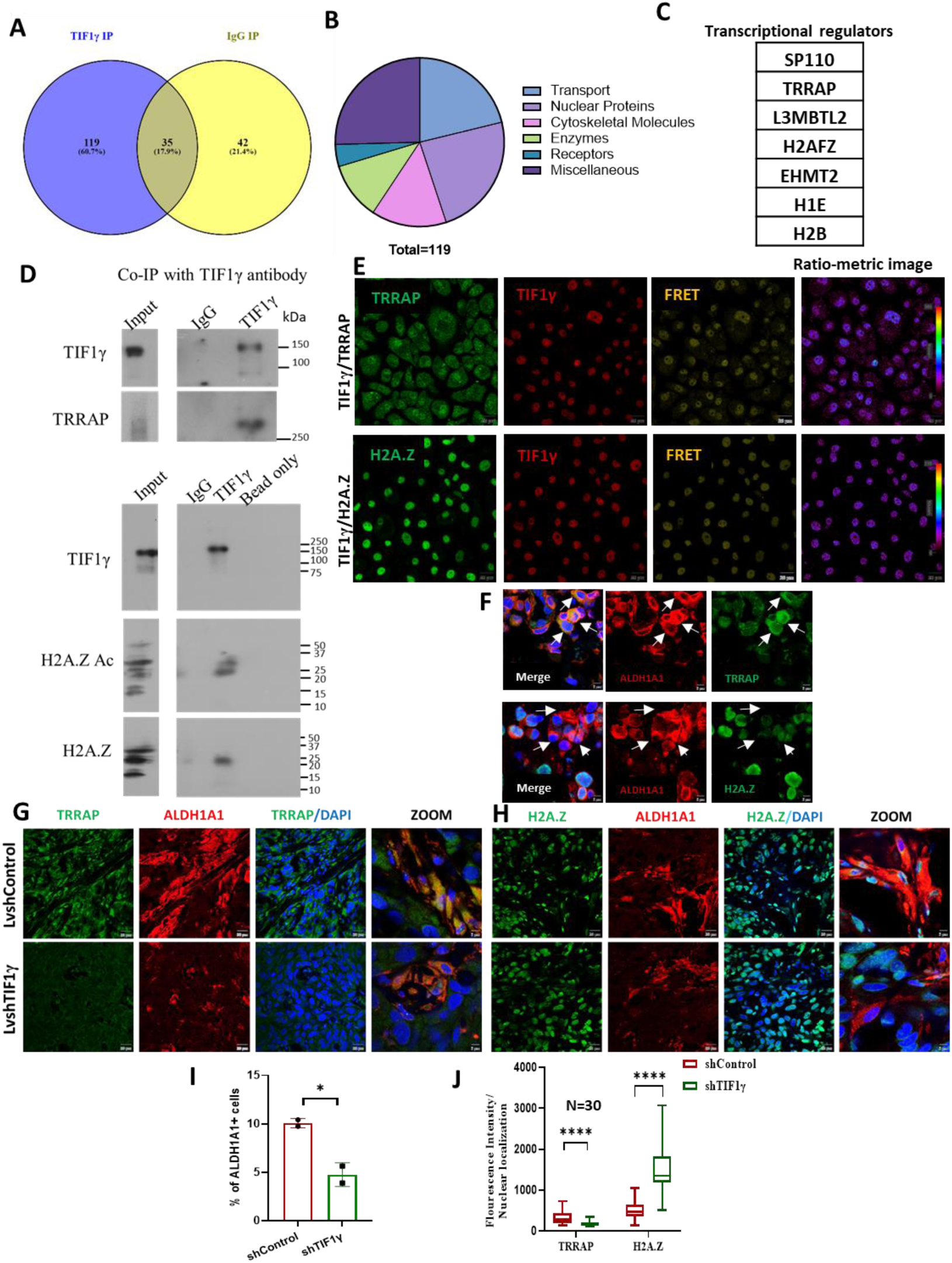
TIF1γ interacts with TRRAP and H2A.Z. (A)The HSC-4 cells were cultured in sphere medium and used for immunoprecipitation (IP) with TIF1γ antibody. The eluate was resolved on SDS-PAGE was given for LC-MS/MS analysis to identify the interacting partners of TIF1γ. The Venn diagram shows the number of molecules identified by LC/MS/MS analysis. (B) Pie-chart of TIF1γ interacting partners based on their function. (C) The set of transcriptional regulators that are TIF1γ interacting molecules. (D) HSC-3 cells were grown for 6 days in sphere culture with MG-132 treatment and were used for the Co-IP experiments. Co-IP was performed with rabbit Anti-TIF1γ or Rabbit IgG (0.5μg) along with 10% input. The eluate was probed for the indicated molecules. (E) The HSC-4 cells were treated with sphere medium for 6 days followed by MG-132 treatment. The secondary antibodies used for immunofluorescence were selected as FRET pairs – Anti-Mouse Alexa 555 was used to stain TIF1γ and Anti-Rabbit Alexa 488 was used to stain TRRAP/H2A.Z. For the FRET imaging excitation was with 488nm and the emission was collected at 595±25. The ratio-metric image generated by cellSens Dimension is shown. (F) Immunohistochemistry on OSCC sections. One set of OSCC sections were stained for TRRAP and ALDH1A1 and the other was stained for H2A.Z and ALDH1A1. The representative confocal image of TRRAP and H2A.Z along with ALDH1A1 in OSCC sections is given. The marked cells represent ALDH1A1 positive CSCs. The scale bar represents 5μm. (G, H) Immunohistochemistry on orthotopic tumour section of both control and TIF1γ knockdown cells. The sections were stained for TRRAP and ALDH1A1 or H2A.Z and ALDH1A1. The scale bar represents 20μm. (I)The quantification of ALDH1A1+ cells in the LvShControl and LvShTIF1γ tumor sections from 60X fields is represented by bar graph. Error bar represents the SD.(J)The fluorescent intensity of TRRAP and H2A.Z in random nuclei is measured and the median expression is plotted. Error bar represents the SD.

A set of 119 molecules were identified to be exclusive for the TIF1γ Co-IP (Figure 4A). The molecules were grouped to those involved in transport, nuclear proteins, cytoskeletal molecules, enzymes, receptors, and the rest were grouped under miscellaneous (Figure 4B). A set of transcriptional regulators having a role in cancer was shortlisted (Figure 4C). Among these molecules, interaction of TRRAP and H2A.Z are reported in Notch signaling pathway, an important pathway in maintaining CSCs (36). It is reported that TRRAP along with Tip60 and P400 acetylates H2A.Z, increasing the chromatin occupancy of H2A.ZAc, but reducing its total form, to switch on the transcription of Notch target gene, like *HES1,* which is a critical regulator of stem cells (36). Next, we examined whether TIF1γ interacts with TRRAP and H2A.Z to perform such a transcriptional regulatory role. The interaction of TIF1γ with TRRAP, acetylated H2A.Z and H2A.Z was confirmed by IP and in the nuclear extracts of sphere cells in presence of proteasomal inhibitor, MG-132 (Figure 4D & S10). Later, we did a FRET analysis to confirm the physical interaction of the molecules using Alexa Fluor 488 and Alexa Fluor 555 as FRET pairs. Although all the cells expressed all the three molecules, the interaction of TIF1γ with TRRAP or H2A.Z was observed in a subset of cells as revealed by the ratio-metric image (Figure 4E). Moreover, the interaction between TRRAP and TIF1γ was stronger than that of TIF1γ and H2A.Z. So, our results revealed a heterogeneity in the cells with respect to the interaction of TIF1γ with TRRAP and H2A.Z.

To further explore this heterogeneity, we examined the spatial distribution of TRRAP and H2A.Z in relation to ALDH1A1 expressing CSCs on primary OSCC sections. The ALDH1A1 high expressing cells, denoting the self-renewing cells, showed high nuclear expression of TRRAP with low H2A.Z nuclear localization (Figure 4F). This negative correlation corroborates with our hypothesis that TRRAP might be acetylating H2A.Z leading to decreased chromatin occupancy. Further the expression of TRRAP and H2A.Z was checked on tumor sections from TIF1γ-depleted and control group along with a CSC marker ALDH1A1 (Figure 4H&I). Parallel to the reduction in ALDH1A1+ve CSC population, there was a decrease in the TRRAP expression with a concomitant increase in the H2A.Z nuclear expression on the TIF1γ-knocked-down tissue sections (Figure 4J). The negative correlation of TRRAP and H2A.Z was evident in our analysis (Figure 4K). Thus, depletion of TIF1γ results in the reduced nuclear localization of TRRAP and increased H2A.Z chromatin occupancy, leading to reduced CSC frequency.

### Functional interaction of TIF1γ/TRRAP/H2A.Z regulate noncanonical *HES1* expression

TRRAP is reported to be a part of acetylation complex involving Tip60 and P400, which acetylates H2A.Z on the promoter, inducing Notch target genes, like *HES1* (36), critically regulating neural stem cell maintenance. In development, Notch-independent expression of *HES1* (Canonical expression) through self-renewal pathways like WNT and BMP pathways support quiescent neural stem cells, while its Notch-dependent expression (canonical expression) leads to the differentiation and proliferation of progenitor population (37). In this context, we evaluated the role of TIF1γ in the regulation of HES1 in oral cancer cells. We observed an enhanced expression of *HES1* upon enrichment of CSCs, in the sphere condition, which was abolished by the depletion of TIF1γ (Figure 5A-B). Since Notch-independent HES1 and Notch-dependent HES1 exerts differential effects, next we checked this aspect using a dual reporter for differential *HES1* expression, which we published earlier (38, 39) (Figure S10A). In the construct, the red expression reports Notch-dependent *HES1* (*NDHES1*) expression (wild type Notch promoter), while the green fluorescent represents Notch-independent *HES1* (*NIHES1*) expression (Notch promoter with mutated RBP-Jκ binding sites).

**Figure 5:**
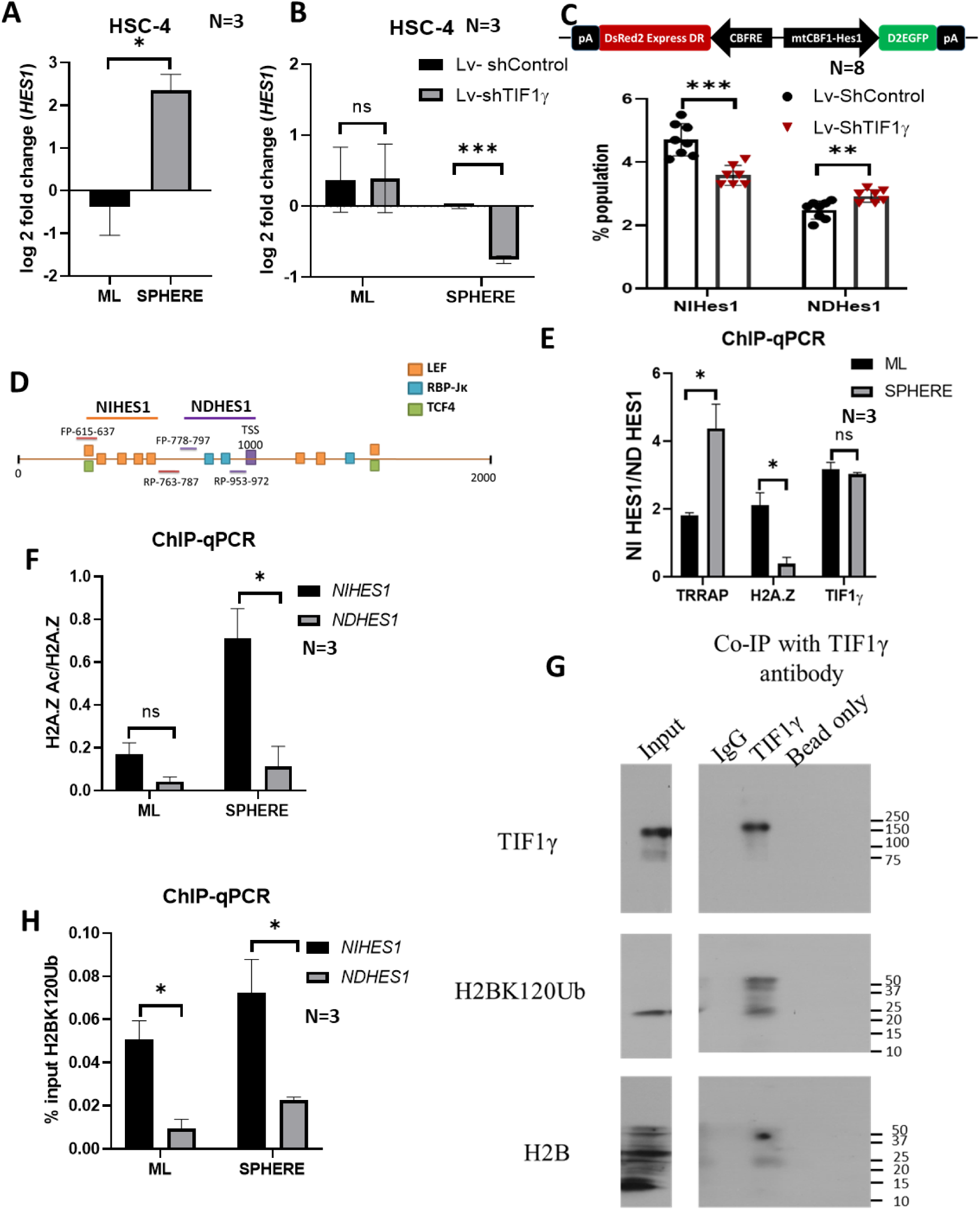
TIF1γ/TRRAP/H2A.Z regulates *HES1* expression. **(A, B)** The HSC-4 cells as indicated were cultured in monolayer and sphere conditions for 6 days and qRT-PCR was performed for *HES1* expression. *β ACTIN* was used as an endogenous control. Error bar represents the SEM.(C) We used a dual reporter were DsRed2 represents NDHES1 and D2EGFP denotes NIHES1. IMR-32 cells harbouring this dual *HES1* reporter with either Lv-Sh TIF1γ or Lv-Sh Control was taken. FACS analysis was done and the data is plotted as mean± SD.(D) Diagrammatic representation of *HES1* promoter denoting the binding sites for RBP-Jκ, LEF and TCF4 identified using TRANSFAC. The primer pairs designed are also represented. (E) The HSC-3 cells were cultured as monolayer and sphere condition for 6 days. A ChIP assay was performed as described under materials and methods. Percentage input was calculated and the graph represents the chromatin occupancy of TRRAP, H2A.Z and TIF1γ on the *NIHES1* compared to *NDHES1*. Error bar represents the SEM.(F) The graph represents chromatin occupancy of H2A.Z Ac compared to total H2A.Z on the *NIHES1* and *NDHES1* promoter in monolayer and sphere condition. Error bar represents the SEM.(G) HSC-3 cells were grown for 6 days in sphere culture with MG-132 treatment and were used for the Co-IP experiments. Co-IP was performed with rabbit Anti-TIF1γ or rabbit IgG (0.5μg) along with 10% input. The eluate was probed for the indicated molecules. (H) A ChIP assay was performed with H2BK120Ub antibody. The graph represents chromatin occupancy of H2BK120ub on the *NIHES1* and *NDHES1* promoter in monolayer and sphere condition. Error bar represents the SEM.

When we used this reporter system to analyze the expression of NIHes1 expressing population (NIHES1) and NDHes1 expressing population (NDHES1), we found that the *HES1* expression observed is mainly by the NIHES1 population, both in monolayer and spheres (Figure S11A). Further, there was a marked increase in the NIHES1 population upon self-renewal enrichment. Moreover, under differentiating condition, the cells harboring this construct showed a significant reduction in NIHES1 with simultaneous increase in NDHES1, upon TIF1γ knockdown (Figure 5C, Figure S10B). So, these results suggested that TIF1γ is required for the induction of Notch-independent *HES1* expression, which is regulated by pathways like, WNT, BMP, TGF-β, etc. So, we next checked if TIF1γ is indispensable for the induction of ALDH1A1 expressing population in response to these signals. Our results showed that abrogation of TIF1γ significantly reduces the induction of CSCs in response to TGF-β1, WNT1 and BMP-4 (Figure S11C). Collectively, our results suggest that TIF1γ might have a transcriptional regulatory role for the expression of self-renewal genes, like *HES1*, specifically through the non-canonical pathway.

To further explore the transcriptional regulation of *HES1* by TIF1γ at the chromatin level, H3K4 trimethylation (H3K4me3) activation mark and H3K9 trimethylation (H3K9me3) repression mark was checked by chromatin immunoprecipitation (ChIP). We designed two primers for ChIP, one for the NDHES1 and the other for NIHES1 regions. One set of primer was designed flanking the RBP-Jκ binding domain representing the NDHES1 expression and the one flanking the LEF binding domain represented NIHES1 expression (Figure 5D). Although there was a preferential expression of NIHES1 over NDHES1, we did not observe any significant difference in the H3K4me3 mark between *NIHES1* and *NDHES1*regions in either monolayer or sphere (Figure S12A, B). The ChIP data of H3K4Me, H3K9Me3 or their relative expression (H3K4Me/H3K9Me3) could not explain the preferential expression of NIHES1 in monolayers or spheres. So, next we explored the possibility of TIF1γ and/or TRRAP regulating chromatin architecture by acetylation for the activation of transcription. First, we tried the component of acetylation complex, TRRAP which associates with TIF1γ and H2A.Z. Our results showed that both in sphere condition and monolayer condition, there was an increased occupancy of TRRAP on *NIHES1* promoter region compared to *NDHES1*. Of note, this enrichment of TRRAP on the *NIHES1* region was significantly more in sphere condition, where self-renewing population is enriched, compared to monolayer cells (Figure 5E). Conversely, H2A.Z occupancy was significantly less on the *NIHES1* promoter region in sphere than monolayer (Figure 5E). When we checked the H2A.ZAc on the promoters, though we observed a trend of preferential occupancy on *NIHES1* region, the difference was not significant. But when we checked the preferential enrichment of H2A.ZAc over total H2A.Z, we observed that in sphere condition, there is a significant enhanced occupancy of H2A.ZAc on *NIHES1* than *NDHES1* (Figure 5F). These results were in agreement with our hypothesis and immunohistochemistry data, which showed that the nuclear localization of TRRAP increases with a decrease in the H2A.Z nuclear localization in CSCs (Figure 4F, H, I). Another important observation was that the occupancy of TIF1γ on *NIHES1* did not change much between monolayer and sphere, questioning the role of TIF1γ in the transcription complex (Figure 5E). Since TIF1γ is reported as a reader for acetylation mark (40), we hypothesized that TIF1γ might be acting as an acetylation reader for H2A.ZAc mark, monoubiquitinating H2B to induce *HES1* expression. We confirmed the interaction of TIF1γ with monoubiquitinated H2BK120 (H2BK120Ub) (Figure 5G, S13). In addition, the occupancy of H2BK120Ub was significantly increased in spheres compared to monolayer condition (Figure 5H). Thus, these results show that TRRAP acetylates H2A.Z and TIF1γ reads this H2A.Z acetylation to monoubiquitinate H2BK120, inducing Notch-independent *HES1* expression.

### TIF1γ transcriptionally regulates self-renewal pathways and genes

Since TIF1γ regulates self-renewal, *HES1* might not be the only self- renewal gene regulated by TIF1γ. In order to identify the TIF1γ target genes under self-renewal condition, an RNA Seq analysis was done to identify the differentially expressed genes in spheres of TIF1γ-depleted cells compared to control spheres (Figure 6A). Our analysis identified 1241 genes (Cut-off: adj.p-value > 0.05, logFC > |2|; 430 down-regulated and 811 upregulated), regulated specifically by TIF1γ, under self-renewing conditions. The volcano plot represents the cut off and the set of down-regulated and up-regulated genes on comparing the TIF1γ knock down with control (Figure 6B).

**Figure 6:**
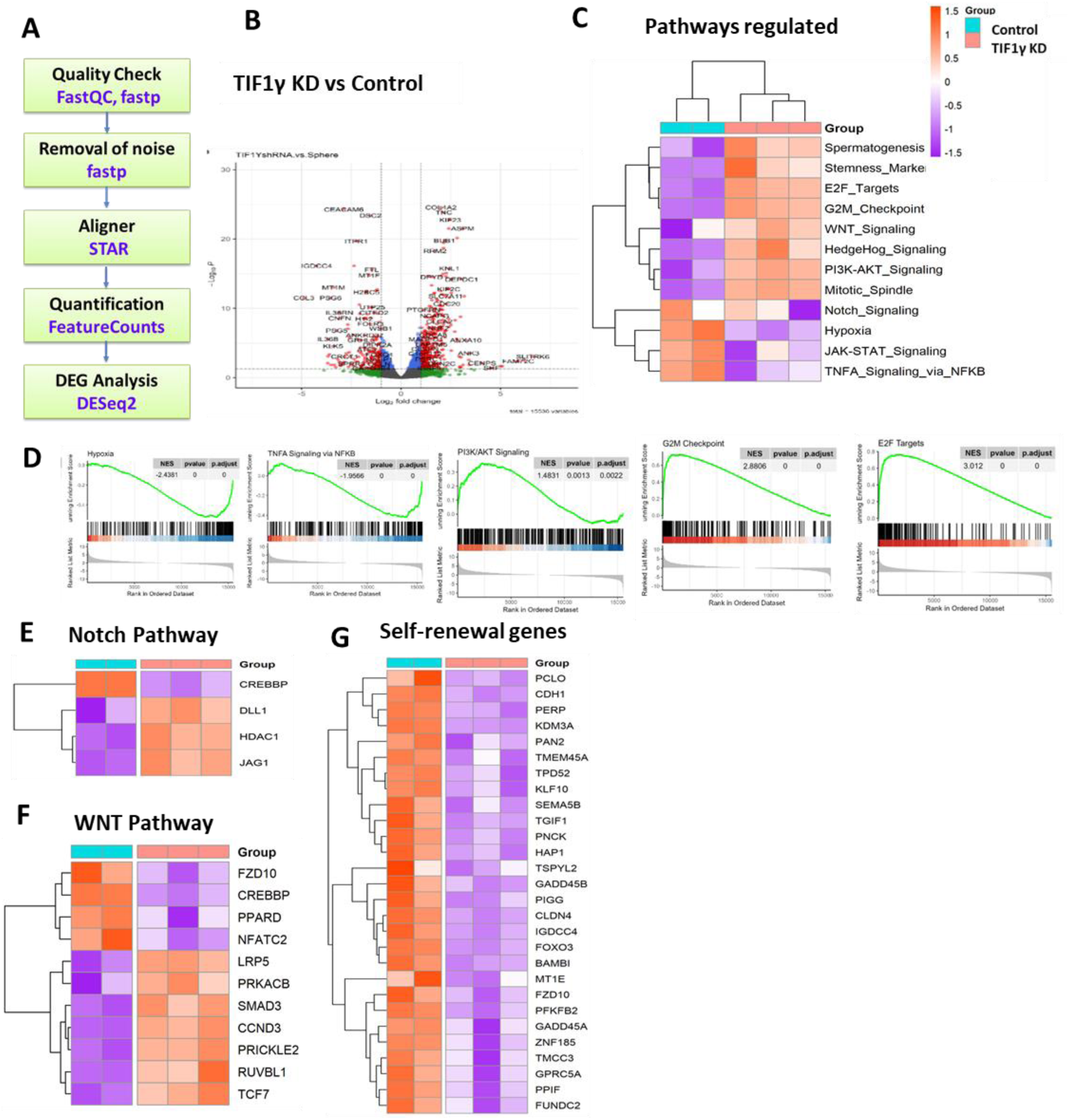
Self-renewal genes regulated by TIF1y (A)HSC-4 cells, either wild type or TIF 1y depleted were cultured in sphere medium for 6 days and RNA Seq was performed. The flow chart for the analysis of the RNA Seq data is given. (B) A Volcano plot showing the differential gene expression analysis between TIF1y depleted and control cells. The threshold values are shown in dashed line (adjusted <0.05 and fold change>1.5). (C) The differential expression data was used for pathway enrichment analysis. Among the pathways, the ones regulating self-renewal was seclected for generating the Heatmap. (D)Among the pathways regulated by TIF1y, Hypoxia pathway and the TNF-u pathway were significantly down-regulated whereas E2F-targets, G2/M checkpomt and PI3/AKT pathway were significantly up-regulated upon TIF1y knockdown. The GSEA plot for these pathways are shown.(E.F) Heatmap for genes significantly regulated under the Notch signalling pathway or WNT pathway upon TIF1y depletion. (G) The orthologues for genes differentially regulated in mouse embryonic stem cells were taken and compared with TIF1y specific genes to obtain TIF1y specific self-renewal genes.

When we analyzed the pathways regulated by TIF1γ, Hypoxia pathway, and the TNF-α pathway were significantly down-regulated whereas E2F-targets, G2/M checkpoint and PI3/AKT pathway were significantly up-regulated upon TIF1γ knockdown (Figure 6C, D). Since the Notch pathway and WNT pathway showed a differential regulation (Figure 6C, S14A), we analyzed the pathway genes in detail. Among the Notch pathway genes, *DLL1* and *JAG1*, the ligands involved in the signaling pathway were up-regulated upon TIF1γ depletion, suggesting that loss of TIF1γ shifts the non-canonical Notch signaling pathway to the canonical one (Figure S14A, 6D). To further ascertain the role of TIF1γ in the maintenance of NIHES1 population, we compared our RNA Seq data with an RNA Seq data we published using neuroblastoma cells. Among the NIHES1-specific genes, a subset of 100 genes showed the reversal of their expression upon the knockdown of TIF1γ, in consistent with our FACS data using the dual HES1 reporter (Figure S14B, 5C). Similar to Notch signaling pathway, TIF1γ loss down-regulated the non-canonical WNT mediator like *NFATC2*, with a simultaneous up-regulation of *TCF7* and *LRP5* that are involved in the canonical WNT signaling (Figure 6F). These data suggest a shift from non-canonical WNT and Notch signaling pathway to the canonical ones upon TIF1γ depletion. To test whether TIF1γ regulates self-renewal genes, we selected a set of self-renewal genes from mouse embryonic stem cells, and their orthologues were compared with TIF1γ specific genes identified in our study. There were 32 embryonic stem cell specific self-renewal genes, whose expressions were significantly diminished with the loss of TIF1γ (Figure 6G).

### TIF1γ/TRRAP/ H2A.Z interaction critically regulates recurrence

Our studies have identified TIF1γ as a regulator of self-renewal. So, we wanted to check the expression of TIF1γ in oral cancer patients. *TRIM33* mRNA expression gradually decreased from normal to dysplasia to OSCC (Figure S15A). When we categorized the OSCC samples based on the grade, the expression significantly increased in the highly aggressive Grade4 compared to Grade3 (Figure S15B). Contrary to the trend observed at mRNA level, we observed that TIF1γ protein expression increased in OSCC and leukoplakia samples compared to normal, as observed by our immunohistochemical analysis (Figure S15C). To test the role of TIF1γ in the prognosis, we checked the correlation to disease free survival. The stage III patient details (N=74) were collected from TCGA data set and the disease-free survival of those patients were analyzed. Consistent with the increase in the highly aggressive form of tumor, the disease-specific survival was high in *TRIM33* low expressing group (Figure 7A). Also, the Kaplan-Meier analysis showed that patients with low expression of *TRIM33* had better prognosis, as revealed by the prolonged disease-free survival or progression-free survival (Figure 7B, C). In conclusion, high TIF1γ expression in advanced stage and grade of oral cancer leads to high recurrence, leading to poor prognosis.

**Figure 7:**
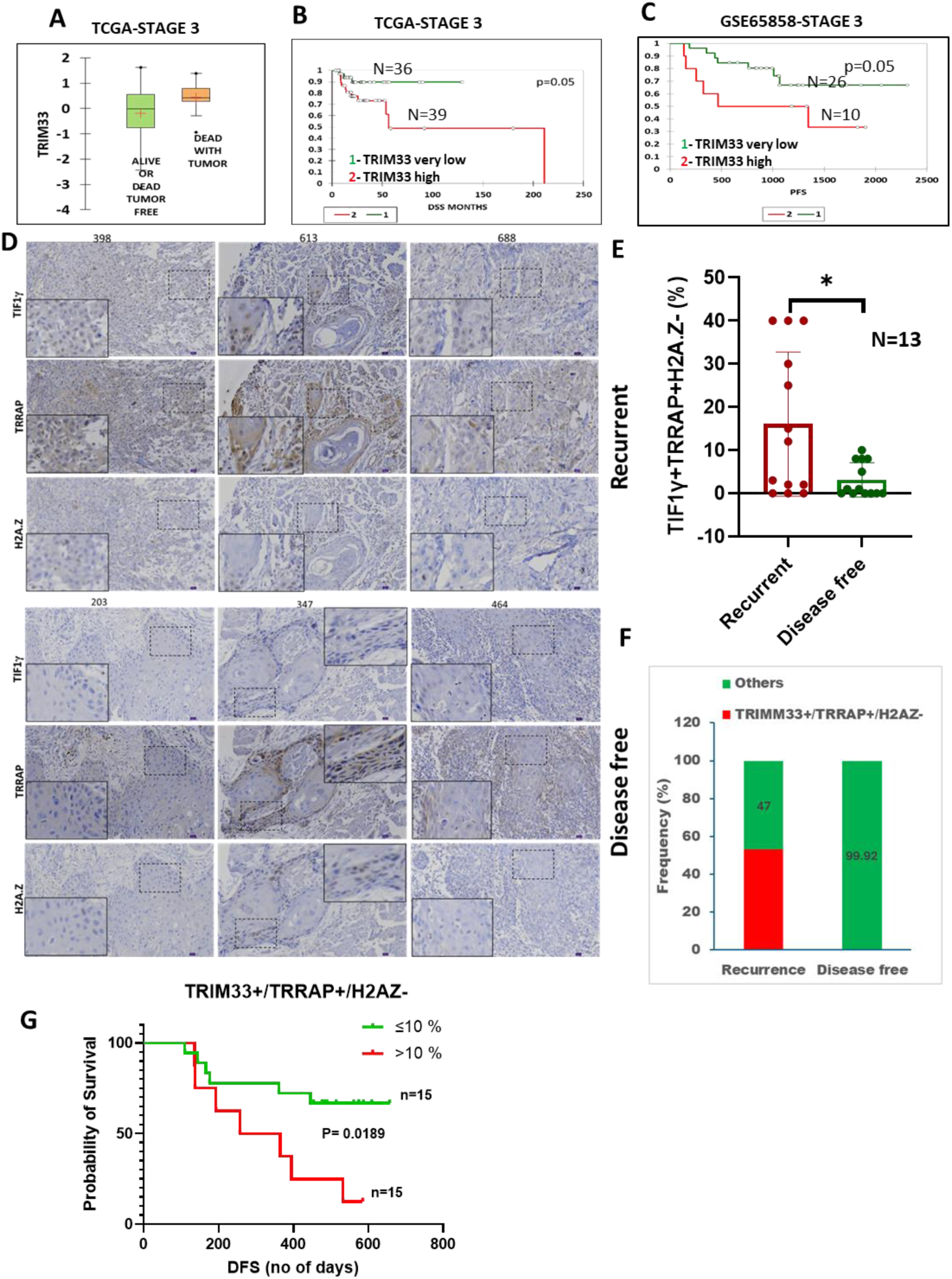
TIF1γ/TRRAP/H2A.Z regulates recurrence. (A)Univariate analysis depicting distribution of TRIM33 levels in oral tumors belonging to stage 3 from TCGA based on the overall survival information. (B)Kaplan-Meier survival analysis in oral tumors belonging to stage 3 from TCGA. (C) Kaplan-Meier survival analysis in oral tumors belonging to stage 3 from GSE65858 dataset. (D) OSCC samples were collected and categorized to recurrent (disease free survival (DFS) less than 15months (N=25)) and disease free (DFS greater than 15months (N=27)). A tissue array was prepared with these samples. Immunohistochemistry was performed with DAB staining. The representative images for staining in recurrent and disease-free group are given. (E) Quantification of TIF1γ TRRAP H2A.Z cells in the recurrence and disease-free group. Error bar represents the SD. (F) The positivity for the three molecules was scored for the same region on the serial section. Number of cells having TIF1γ TRRAP H2A.Z trend were scored. Number of samples with ≥10% were plotted in recurrent (N=13) and disease free (N=13) group. (G) Samples were grouped as <10% or ≥10% of TIF1γ TRRAP H2A.Z positivity and a Kaplan-Meier survival analysis for DFS was plotted.

The earlier finding suggested that TIF1γ is functionally co-operating with TRRAP and H2A.Z and trigger transcription of a set of self-renewal genes, which could be crucial for recurrence. A correlation study was performed using TCGA data set to see if these molecules are functionally cooperating in clinical samples. In accordance with our wet lab results, *TRIM33* and *TRRAP* showed a significant positive correlation while *TRIM33* and *H2AF.Z* exhibited a weaker negative correlation (Figure S16 A). In spite of that, these three genes showed a functional cooperation, reflected by the high average gene score (0.841, p<0.0001) (Figure S16 B). Yet, the expression of Notch target genes *HES1* or *MYC*, were not correlated to *TRIM33, TRRAP* and *H2AF.Z* (Figure S16 B). Interestingly, hierarchical clustering heatmap analysis revealed that even though patients who had recurrence showed higher expression of all the three genes, *TRIM33,* and *TRRAP* were highly linked to recurrence (Figure S16 C).

In order to visualize the functional cooperation of TIF1γ, TRRAP and H2A.Z at cellular level, we performed immunohistochemistry (IHC) using a tissue microarray of OSCC patients who had either recurrence within 15 months or had disease free survival for more than 15 months (Figure 7D). We observed that in the recurrent group majority of the cells that were positively stained for TIF1γ and TRRAP showed negative staining for H2A.Z and the TIF1γ^+^/TRRAP^+^/H2A.Z^-^ cells came up to 60%, while in the non-recurrent group it was less than 10%. In the recurrent group, 53% of the samples showed more than 10% TIF1γ^+^/TRRAP^+^/H2A.Z^-^ positivity, while in the disease-free group there was only one sample that showed this trend, constituting 0.08% (Figure 7E). The Kaplan-Meier plot showed that the TIF1γ^+^/TRRAP^+^/H2A.Z^-^ expression correlates to poor prognosis, as this group exhibited higher chance of recurrence (Figure 7F). These results reinforce the critical transcriptional regulatory role of TIF1γ in oral cancer recurrence.

## DISCUSSION

The deregulation of TGF-β signaling is a crucial step in the tumor progression, as its tumor suppressive role in the initial stages of cancer is replaced by a tumor promoting role by a complex interplay of signaling cross-talk. Since TGF-β by the interaction with its receptor can lead to its effect either by canonical Smad 4 pathway or noncanonical pathway, it is crucial to understand the molecular intermediates for the tumor regulatory functions. As reported by many others in different cancers, we observed that TGF-β signaling is crucial for the self-renewing CSCs of oral cancer, and it can regulate the growth of drug resistant population, which is vital for recurrence (Figure 1A, B) (41–44). An in-depth analysis of the molecular mechanism behind the TGF-β-mediated self-renewal induction in CSCs is not reported. The importance of a noncanonical TGF-β pathway in CSCs, suggested by our phosphoproteomic analysis was substantiated by our extreme limiting dilution assay (ELDA) (Figure 1C-E, S1). Consistent with our observation that Smad 4 is dispensable for the CSC properties, Smad 4 inhibition has been shown to impart self-renewal in melanoma and neuroblastoma cells (45, 46). More importantly, a TGF-β receptor-independent phosphorylation of Smad 2/3 at the linker region which blocks the canonical TGF-β signaling is shown to be a marker of CSCs (47–50). However, Smad 4 and the canonical signaling are well-characterized as protumorigenic (2). Consistent with that we observed a reduction in the tumor proliferation in our 3D culture on alginate matrix, when Smad 4 was depleted (Figure 1C). Though Smad 4 has protumorigenic functions, the induction of CSCs by TGF-β is not dependent on the canonical Smad 4 dependent pathway.

TIF1γ, identified as a noncanonical mediator of TGF-β, is implicated in other self-renewal pathways like WNT (5, 51, 52) and BMP (53). In hematopoiesis, TIF1γ is essential to maintain the pluripotency of LT-HSCs, and its depletion leads to their differentiation to myeloid lineage (13). Consistent with its role in development, our study using *in vitro* and *in vivo* models has shown that TIF1γ is essential for the maintenance of CSCs of oral cancer, as evidenced by our CSC reporter ALDH1A1. ALDH1A1 is a marker of CSCs in a wide variety of cancers (54–59), and its expression is correlated to poor prognosis (60–64). The data we generated by the reporter construct was validated with functional assays like ELDA and other pluripotent markers (Figure 2). Generally, TIF1γ is considered as a tumor suppressor as it inhibits epithelial to mesenchymal transition (EMT) by different ways (5, 20, 21, 65, 66). One of the major means of tumor suppression is the E3 ubiquitin ligase-mediated repression of EMT regulators (20, 66, 67). Here, we identified a novel mode of action of TIF1γ, as a transcriptional regulator of self-renewal genes, supporting cancer progression and poor prognosis. Though TIF1γ is shown to support breast cancer and prostate cancer progression, the mode of action was stabilization of the nuclear receptors, which is quite distinct from the histone acetylation reader activity, we identified in this study (Figure 5)(6, 7).

Though the presence of bromodomain predicts the histone acetylation reader activity for TIF1γ, the evidence for this activity came recently, where it was shown to read the acetylation of histones marked by TRRAP-Tip60 complex on the HSP72 promoter. Upon reading the acetylation mark, TIF1γ monoubiquitinated H2BK120 (H2BK120Ub) residue, which initiates the transcription (40). In our study, TIF1γ associated with TRRAP and Acetylated H2A.Z on the promoter of *HES1*, specifically on the LEF-binding region upon induction of self-renewal, as shown by our IP and ChIP analysis (Figure 4,5). H2A.Z is an evolutionally conserved histone variant, which is functionally different from replication-dependent H2A. It plays a crucial role in varied biological functions, including transcriptional activation and repression, DNA damage recognition and repair, nucleosome stability, chromosome organization as well as stem cell differentiation and maintenance (68). H2A.Z can exert either transcriptional activation or repression, primarily depending on its posttranslational modifications such as acetylation, methylation, ubiquitination and SUMOylation (68). Acetylated H2A.Z (H2A.Z Ac) is shown to occupy the promoters of active genes, suggesting the role of acetylation in the activation of transcription (68–71). ChIP Seq analysis has shown that the occupancy of total H2A.Z on the promoters were more on the repressive chromatin region whereas the H2A.Z Ac occupancy was predominantly on the active and poised chromatin in cancer cells (69). A reverse correlation of H2A.Z occupancy with H2A.Z Ac and active transcription is demonstrated in the transcription of Notch target genes, like *HES1* (36). In agreement to this, we found decreased H2A.Z but increased H2A.ZAc on the NIHES1 region of *HES1* promoter in sphere condition (Figure 5). The reverse correlation of H2A.Z and active transcription of self-renewal genes like *HES1* was evident from our ChIP analysis, which was further reinforced by our immunohistochemical analysis on xenograft sections and OSCC samples (Figure 4F-J, 5F-H, 7D). The importance of H2A.Z occupancy in the repression of poised chromatin is shown in stem cell regulation, supporting our observation that H2A.Z occupancy is considerably less in ALDH1A1expressing CSCs (68, 72, 73). Although TRRAP-Tip60 complex as an acetyl transferase for H2A.Z is well established in the literature, its association with TIF1γ, acting as a reader for the H2A.Z Ac is our novel finding (36, 68, 71). Taken together, our study unraveled a novel protumorigenic role of TIF1γ, as a transcriptional activator of *HES1* by virtue of its histone acetylation reader activity as well as E3 Ubiquitin ligase activity.

*HES1* is a self-renewal gene, reported to be involved in maintaining normal stem cells and CSCs (37, 39, 74–76). Earlier, we have shown that HES1 plays a critical role in neural differentiation during neocortical development (37, 39). With the same dual reporter system that we used in the present study, we showed that NIHes1 expression is limited to quiescent stem cells whereas the expression of NDHes1 is observed in the proliferating radial glial cells, in the progenitor territory (38). In parallel to this, in the human cancer counterpart, the NIHES1 expressing subpopulation represents the more primitive CSCs than the NDHES1 expressing CSCs (39). In the present study, we showed that TIF1γ depletion induces differentiation of NIHES1 population to NDHES1 population (Figure 5C). We have also shown that TIF1γ is critical for the induction of self-renewal by WNT and BMP, two major pathways regulating NIHES1 expression (Figure S11C) (37). Though Notch pathway is a known stem cell and CSC regulator, our previous work has shown that while noncanonical Notch supports the primitive quiescent stem cells, the canonical Notch drives the proliferation of progenitor population or progenitor-like CSCs (38, 39, 77). Even though our RNA Seq results showed that the Notch pathway is down-regulated upon TIF1γ depletion (Figure 6C), the analysis of the target gene list suggested the conversion of Notch-independent state to Notch–dependent state, upon reduction of TIF1γ expression, as evidenced by the upregulation of Notch ligands, *DLL1* and *JAG1* (Figure 6E). Interestingly, we observed a reversal of expression pattern of 100 NIHHES1 specific genes, upon knockdown of TIF1γ in oral cancer cells, further arguing the role of TIF1γ in the expression of NIHES1 specific self-renewal genes (Figure S14B) (39). Apart from Notch signaling, WNT signaling also showed a shift from noncanonical signaling to canonical signaling when TIF1γ was knocked-down (Figure 6F). The noncanonical WNT signaling and its cross talk with Notch signaling is shown to regulate self-renewal of normal stem cells (78–80). As expected, the noncanonical WNT signaling through FZD10 and NFATC2, the WNT regulators we observed to be down-regulated upon TIF1γ depletion, is regulating CSCs and cancer progression (Figure 6E&F) (81–84). Up-regulation of proliferation pathways like E2F, G2/M and mitotic spindle upon TIF1γ knockdown suggest that TIF1γ may be essential to block proliferation to maintain stem cell population. We have also observed the downregulation of a set of pluripotency genes identified in embryonic stem cells, verifying the role of TIF1γ in the maintenance of self-renewal.

Since our functional analysis confirmed that TIF1γ regulates self-renewal of cancer cells by the transcriptional upregulation of a set of self-renewal genes with its histone acetylation reader activity and E3 ubiquitin ligase activity, we tested the impact of this molecule in the disease. Our immunohistochemical analysis showed that there is heterogeneity in the expression of TIF1γ in cancer cells, and it is limited to certain clusters (Figure S15C, Figure 7D). Based on our functional analysis, we hypothesized that in a given tumor microenvironment, the TIF1γ expressing cells might have the capability to sustain self-renewal. Our results confirmed that the TIF1γ^−ve^ population gradually gets depleted in a tumor microenvironment due to the lack of self-renewal ability (Figure 2F-H). Further, we showed that the induction of self-renewal ability is a result of the complex formation of TRRAP and H2A.Z with TIF1γ (Figure 4F-J, 7D). Our speculation that the TIF1γ-mediated sustenance of self-renewal might negatively influence the prognosis of the disease was strengthened by our *in-silico* analysis of the TCGA data set. The Kaplan Meier survival analysis showed that TIF1γ gene *TRIM33* is inversely correlated to disease free survival (Figure 7 A-C). The observation that *TRIM33, TRRAP* and *H2AF.Z* functionally cooperated and the positive correlation of *TRIM33* to *TRRAP* and its negative correlation to *H2AF.Z* reinforced the importance of the novel molecular mechanism we unraveled in cancer progression (Figure S16). We tested whether TIF1γ loss will improve the prognosis of the disease by reducing recurrence, using an orthotopic floor of the mouth oral cancer model. Our observations confirmed that TIF1γ regulates recurrence of the disease and the reduction of TIF1γ expression can improve the prognosis (Figure 3). The significance of our result was validated using primary OSCC samples. Our immunohistochemical analysis of the tissue microarray proved that as the population of cancer cells showing the functional cooperation of TIF1γ, TRRAP and H2A.Z increases, the chance of recurrence of the disease increases.

To support the notion that tumor cells hijack and activate the normal embryonic development pathways, we identified a novel TIF1γ-mediated protumorigenic mechanism in oral cancer. TIF1γ was shown to be critical for the maintenance of pluripotent LT-HSCs during hematopoiesis, and the loss of function of TIF1γ leads to aberrant differentiation. In this paper, we demonstrated that a parallel mechanism exists in oral cancer, and moreover, we unraveled the transcriptional regulatory mechanism by which TIF1γ exerts this self-renewal function. Since E3 ubiquitin ligase activity is shown to be the major role of tumor suppressor function of TIF1γ while acetylation reader activity and E3 ubiquitin ligase activity together impose the tumor promoter function, the acetylation reader activity can be targeted to exclusively block the tumor promoter function. So, we propose that the TIF1γ bromodomain can be a therapeutic target for oral cancer, which needs further validation.

## MATERIALS AND METHODS

### Ethics statement

The studies involving human participants were reviewed and approved by Institute Human Ethical Committee, Rajiv Gandhi Centre for Biotechnology (IHEC/1/2011/04, IHEC/8/2021_2/13). The patients/participants provided their written informed consent to participate in this study. Paraffin embedded tissue blocks for making the tissue microarray (TMA) were collected from the archives of Cochin Cancer Research Centre, Cochin, and the preparation of TMA, IHC staining, scoring and imaging were done at St. Johns Research Institute. The protocol was approved by Institute Human Ethical Committee of all the respective study centers (IEC/150/2023, IHEC/1112022_Exl11). The use of plasmids and lentiviral shRNA were reviewed and approved by Institutional Bio-safety Committee (66/IBSC/TTM/2022). The animal studies were reviewed and approved by Institute Animal Ethics Committee (IAEC/873/TM/2022), Rajiv Gandhi Centre for Biotechnology, Thiruvananthapuram.

### Antibodies

The antibodies, TIF1γ, TRRAP, Acetyl-Histone H2A.Z (Lys4/Lys7), H2A.Z, Ubiquityl-Histone H2B (LyS110), H2B, Protein A HRP and rabbit IgG were purchased from Cell Signaling Technology, Danvers,755 Massachusetts, USA. While the other antibodies, Mouse TIF1γ, EpCAM, TGF-β, Smad4, β-Actin and mouse IgG were purchased from Santa Cruz Biotechnology, Inc., Dallas,754 Texas, USA. Peroxidase AffiniPure Donkey Anti Mouse and Peroxidase AffiniPure Donkey Anti Rabbit were purchased from Jackson ImmunoResearch Laboratories Inc., West Baltimore Pike, West Grove, PA 19390, USA. The IP detection reagent, Horseradish Peroxidase conjugated Veriblot was purchased from Abcam, Cambridge, United Kingdom. The Donkey Anti-Rabbit Alexa Fluor-488, Donkey Anti-Mouse Alexa Fluor-488, Donkey Anti-Rabbit Alexa Fluor-568, Donkey Anti-mouse Alexa Fluor-568, Donkey Anti-Rabbit Alexa Fluor-680, Donkey Anti-mouse Alexa Fluor-680 and Donkey Anti-mouse Alexa Fluor-555 from ThermoFisher Scientific, Waltham, Massachusetts, USA were used for FRET, immunofluorescence and immunohistochemistry experiments.

### Plasmids and shRNAs

DsRed2 N1 was a gift from Michael Davidson (Addgene plasmid 54493). pRetroSuper-GFP TIF1gamma was a gift from Joan Massague (Addgene plasmid 1572). pCBFREDsRed-Express-DR-mtCBF1-Hes1-d2EGFP, the dual reporter for Hes-1 promoter was a kind gift from Dr. Jackson James, Rajiv Gandhi Centre for Biotechnology, Thiruvananthapuram. Lentiviral shRNAs – LvshControl (sc-44231-V) and LvshTIF1γ (sc-63127-V) were purchased from Santa Cruz Biotechnology, Inc., Dallas,754 Texas, USA.

### Cell lines

The cell lines RCB1015, RCB1017, HSC-3 and HSC-4 were obtained from Riken Cell Bank, Japan. All these cell lines were maintained in Dulbecco’s Modified Eagle’s Media supplemented with 10% FBS. HSC-3 and HSC-4 cells stably expressing ALDH1A1-DsRed2 were maintained in 50µg/mL geneticin containing medium. Cells containing the shRNA were maintained in 100ng/mL puromycin containing medium.

### Monolayer and Sphere Culture

The Sphere culture is established with sphere medium which is constituted by 1X OptiMEM, reduced sphere medium, supplemented with 1X Insulin- Transferrin- Selenium, 1X N2, 20μg/ml EGF and bFGF. on a non-adherent dish (coated with 1% Poly (2-hydroxyethyl methacrylate) or polyHEMA to deplete adherence of the dish). Monolayer culture was maintained in DMEM supplemented with 10% FBS. All required reagents were purchased from ThermoFisher Scientific, Waltham, Massachusetts, USA.

### Lentiviral Infection

The lentiviral infection of shRNA was done in BSL-II facility. The cells were seeded at a density of 0.15×10^6^ cells in a 24 well plate. Next day, 20μg/mL sequabrene was added to the cells in 250μL10%DMEM. 2.5µL of lentiviral particle was added from the stock and mixed. A medium change was given after 24hrs. After 48hours, the cells were trypsinized and seeded for selection.

### Serial dilution sphere formation assay

Different dilutions of cells were seeded in sphere medium in 96 well plate coated with 1% polyHEMA by serial dilution. Each dilution had 8 replicates in one 96 well plate. The dilution for each cell line was standardised separately. The cells were grown in sphere medium for 6days with medium supplementation every third day, and on the 6^th^ day the number of spheres in each well was counted.

### ELDA analysis

ELDA analysis to calculate the stem cell frequency was done online using ELDA webtool (https://bioinf.wehi.edu.au/software/elda/) designed by the Bioinformatics Division, The Walter and Eliza Hall Institute of Medical Research in Australia. The stem cell frequency is calculated by slope of the line graph plotting log fraction of non-responders and dilution factor.

### RNA isolation and qRT-PCR

RNA was prepared using RNeasy Mini Kit as per manufacturers instruction. RNA was reverse transcribed using Maxima reverse transcriptase and Ribolock RNase inhibitor Kits. The gene expression was checked using PCR with gene specific primers which was run on agarose gel for electrophoresis. The bands were visualized in Gel Doc EZ imager (Bio-Rad Laboratories, Hercules, California, USA). Quantification of gene expression and ChIP samples were done by qRT-PCR using Power SYBR Green Master Mix and analyzed using Quantstudio^TM^ PCR system (Applied Biosystems). Data for gene expression was normalized using β Actin as the house keeping gene. All the reagents for cDNA synthesis, PCR and qRT-PCR were purchased from ThermoFisher Scientific, Waltham, Massachusetts, USA while the RNeasy Mini Kit which was purchased from QIAGEN, Hilden, Germany.

Primers for *NANOG* are F: 5’- TGCAGAGAAGAGTGTCGCA -3’, R: 5’-GGTCTTCACCTGTTTGTAGCTG -3’. Primers for *BMI 1* are F: 5’-TTCTGCTGATGCTGCCAATG -3’, R: 5’- TCCGATCCAATCTGTTCTGGT -3’. Primers for *KLF4* are F: 5’- TGCCCCGAATAACCGCT -3’, R: 5’- CGTTGAACTCCTCGGTCTCT -3’. Primers for *OCT4A* are F: 5’- TCCAGGTGGTGGAGGTGAT -3’, R: 5’-CCCCCACAGAACTCATACGG -3’. Primers for *SOX2* are F: 5’-CGCCGAGTGGAAACTTTTGTC -3’, R: 5’- CGCTCGCCATGCTATTGCC -3’. Primers for *Hes1* Notch independent promoter are F: 5’- ACTTACTACAGTCAAAGCAGCTC-3’, R: 5’-AACGTCAATCAAAAGGATTTTAACC-3’. Primers for *Hes1* Notch dependent promoter are F: 5’- GATTGACGTTGTAGCCTCCG-3’, R: 5’- ATATCTGGGACTGCACGCGA-3’. Primers for *HES1* are F: 5’- GATTGACGTTGTAGCCTCCG-3’, R: 5’- ATATCTGGGACTGCACGCGA-3’. Primers for *β-Actin* are F: 5’-CCTTCCTTCCTGGGCATGG-3’, R: 5’-CGCTCAGGAGGAGCAATGA-3’.

### Transfection

Cells were seeded at a density of 0.6x10^6^. Next day, 1.2μg of DNA and 3 μl Lipofectamine 2000 each were diluted in 60μl OptiMEM and incubated at room temperature for10 minutes. The diluted DNA was mixed with the diluted Lipofectamine 2000 and incubated at room temperature for 40minutes. Simultaneously the cells were starved with OptiMEM for 30 minutes. The DNA-Lipofectamine mix was added to the cells such that the final volume with OptiMEM was 1.2ml. Cells were incubated for 4hours in incubator followed by a medium change with OptiMEM and incubation for 30 minutes. The cells were supplemented with 10%DMEM and expression was checked after 48 hours.

### Cell lysis

Total cell lysate was prepared by using lysis buffer (Tris 25mM, NaCl 150mM, EDTA 1mM, NP-40 0.5%, Triton X-100 0.5% and Glycerol 5%) followed by 5 times freeze (liquid N2)-thaw (37°C) cycle and syringe homogenization. The protein was quantified by DC protein assay.

### Nuclear extraction and Immunoprecipitation

The cells were cultured in sphere medium for 6 days. On the 6^th^ day, cells were treated with MG-132 (50nM) for 5hours. The nuclear extraction buffer (0.1mM EGTA, 0.1mM EDTA, 10mM HEPES pH7.9, 10mM KCl and 1mM DTT) was added to the cells and incubated on ice for 20 minutes. The cells were scraped and centrifuged. The pellet was resuspended in lysis buffer (Tris 25mM, NaCl 150mM, EDTA 1mM, NP-40 0.5%, Triton X-100 0.5% and Glycerol 5%) followed by 5 times freeze (liquid N2)-thaw (37°C) cycle and syringe homogenization. The protein was quantified by DC protein assay.

1mg of protein was taken in 500µL for pre-clearing. Simultaneously antibody was conjugated with protein-A/G dynabeads by incubation for 4 hours at 4°C with mixing. The pre-cleared lysate was added to antibody-bead conjugate and incubated overnight at 4°C. Next day, the complex was washed with PBS and denaturing elution was done with 2X loading dye and run on an SDS-PAGE and blotted onto nitrocellulose membrane to probe with specific antibodies.

### Western blotting

A minimum of 40ug protein is loaded onto gel during SDS-PAGE. The proteins were then transferred to nitrocellulose membrane. After a 10 minutes wash with TBST buffer, membrane was blocked with corresponding blocking buffer (10% milk, 5% milk or 3% BSA in TBST) for 1 hour and the blots were incubated with primary antibody overnight at 4°C overnight. Next day, the blots were washed thrice in TBST and incubated in HRP-conjugated secondary antibody for 1 hour at room temperature. The blots were again washed thrice with TBST and stored in TBS until developing. The substrate for HRP, peroxide and luminol from Clarity Western ECL Substrate from BioRad was mixed in equal proportion and added onto the blot. The bands were obtained on an X-Ray sheet after giving proper exposure. The blots were scanned to the computer. The bands were quantified in terms of integrated density in the ImageJ software. The normalized integrated density by the β- Actin bands were plotted in the graph.

### Immunoprecipitation and LC-MS/MS analysis

The cells were lysed and protein was quantified. 1mg of protein was taken in 500µL for pre-clearing. Simultaneously antibody was conjugated with protein-A/G dynabeads by incubation for 4 hours at 4°C with mixing. The pre-cleared lysate was added to antibody-bead conjugate and incubated overnight at 4°C. Next day, competitive elution with antibody epitope peptide was done to elute the protein-protein conjugate from dynabeads. This eluate was resolved on an SDS PAGE for 1cm and the gel pieces were given for Liquid Chromatography- Mass Spectrometry with two detectors (LC-MS/MS) analysis, **Thermo LTQ Orbitrap Discovery in C-CAMP, NCBS** to identify the interacting partners of TIF1γ.

### Chromatin immunoprecipitation

2.5 million cells were taken. Cells were fixed with 1%Formaldehyde followed by quenching with 150mM Glycine. The cells were washed with PBS twice and resuspended in Lysis buffer. The cells were sonicated and the sample was diluted with ChIP dilution buffer. The sonication of sample was confirmed in agarose gel. Input was collected at this step and rest of the sample was incubated with antibody at 4°C overnight. The overnight sample was incubated with Protein A/G Dynabeads for 2 hours at 4°C. The bound complex was washed with ice cold RIPA 150 wash buffer, RIPA 500 wash buffer, LiCl wash buffer and 10mM Tris-Cl pH 8.5 sequentially. The antibody- chromatin complex was eluted with elution buffer (0.5 M Sodium bicarbonate+1% SDS and 5mM DTT). The eluate was treated with RNase A (0.005 ug/ul ) for 30 minutes at 37°C and Proteinase K (0.6 ug/ul) for 3hours at 65°C. The eluate and input were then purified with PCR and Gel clean up Kit.

For designing ChIP primers, the binding domain of RBP-Jκ (for Notch binding), TCF4 and LEF (WNT transcription factors) was identified using TransFac software. RBP-Jκ is involved in canonical notch signaling pathway and hence will represent the Notch dependent HES1expression. The LEF-1 and TCF-4 are WNT signaling molecule which will represent Notch Independent HES1 expression. The RBP-Jκ binding domains were identified at 912-923, 936-947 and 1433-1444 base pairs. The LEF-1 binding domains were found at 623-630, 662-669, 702-709, 733-740, 754-761, 1312-1319, 1362-1369, 1470-1477 and TCF-4 binding domains were found at 622-631, 1469-1478 base pairs. One primer set was designed flanking RBP-Jκ binding domain which will represent *NDHES1* promoter region while the other primer set was designed flanking LEF-1 binding domain which will represent *NIHES1* promoter region. Primers for *Hes1* Notch independent promoter are F: 5’-ACTTACTACAGTCAAAGCAGCTC-3’, R: 5’- AACGTCAATCAAAAGGATTTTAACC-3’.

Primers for *Hes1* Notch dependent promoter are F: 5’- GATTGACGTTGTAGCCTCCG-3’, R: 5’- ATATCTGGGACTGCACGCGA-3’.

qPCR results obtained as cycle threshold (Ct) values. The gene expression of specific gene was normalized with expression of *β-Actin* followed with fold change calculation using 2^-ΔΔCt method. For ChIP- qPCR, the adjusted input Ct (Input Ct- dilution factor) value is subtracted from immunoprecipitated Ct value to obtain ΔCt. The percentage input is calculated by 100/2^ΔCt which represents the enrichment of the target DNA in the IP sample relative to the starting material.

### *In vivo* serial dilution xenograft assay

The cells were trypsinized and counted to determine the cell density. All the cell lines were serially diluted to 10^6^,10^5^ and 10^4^ numbers and injected each dilution into 5 NOD.Cg-Prkdc^scid^/J (NOD-SCID) or NOD.Cg-Prkdc^scid^ Il2rg^tm1Wjl^/SzJ (NSG) mice with 50μL matrigel, subcutaneously on the flanks for xenograft generation. After 21 days, the animals were euthanized, and tumors were collected. The weight of the tumor was determined to plot the graph.

### Orthotopic model for recurrence

HSC-4 cells with stable luciferase expression had been taken and TIF1γ was down-regulated in these cells. These cells were injected orthotopically into the NOD.Cg-Prkdc^scid^ Il2rg^tm1Wjl^/SzJ (NSG) mice under the tongue below mylohyoid muscles with 50µL matrigel. 10 animals were injected in each control and TIF1γ knockdown group. The growth of the tumor was tracked by bio-imaging. Animals were imaged using Perkin Elmer IVIS SPECTRUM *in vivo* imaging system under 2% isoflurane anesthesia. For bioluminescence imaging, the substrate D-luciferin dissolved in saline was injected intraperitonially (80mg/ Kg body weight) and imaging was done after 10 minutes. Once the tumor was grown to 0.5cm^3^ in size, the resection surgery was done. The complete removal of the tumor was confirmed by bio-imaging. The animal was further given post-operative care and imaged periodically for checking the relapse.

### Immunofluorescence and Immunohistochemistry staining

Oral cancer cells were seeded in confocal dish. The cells were washed with PBS and fixed with 4% PFA for 10 minutes. PFA was washed off with PBS, thrice. Cells were permeabilized and blocked by incubating with 2%FBS in PBST (PBS+0.3% Triton X-100) for 50 minutes to 1hour. After blocking. the cells were incubated overnight at 4°C with Primary antibody diluted in 2% FBS in PBS. The following day, the dishes were washed thrice with PBS. Secondary antibody conjugated with fluorophore diluted in 2% FBS in PBS was added to the cells and incubated for 1 hour at room temperature. The nuclei were stained with DAPI followed by three PBS washes to removal of non-specific binding. cells were mounted with ProLong™ Gold Antifade Mountant (ThermoFisherScientific, Waltham, Massachusetts, USA) and imaged with a confocal microscope, Olympus FV3000 or Nikon A1R LSCM.

Tumor samples collected from animal experiments were fixed with 4% PFA for 1 day, followed by storage in 30% sucrose solution. Then, 5μm sections were taken using the LeicaCM1850UV Cryostat. The antigen unmasking was done using 0.01M Citrate buffer, pH 6.0. The immunohistochemistry staining protocol was similar to immunofluorescence staining.

### Fluorescence Resonance Energy Transfer (FRET)

The secondary antibodies used for immunofluorescence were selected as FRET pairs – Anti-Mouse Alexa 555 was used to stain TIF1γ and Anti-Rabbit Alexa 488 was used to stain TRRAP/H2A.Z. Controls with each molecule stained alone were also kept to nullify the spectral overlap. The imaging was done with Olympus FV3000 confocal microscope in 60X magnification. Three different images were taken in three different channels. TIF1γ was imaged by exciting Alexa 555 fluorophore and collecting the emission in the corresponding wavelength detector. TRRAP/H2A.Z was imaged by exciting Alexa 488 fluorophore and collecting the emission in the corresponding wavelength detector. The FRET channel image was obtained by exciting the donor Alexa 488 and collecting the emission from the acceptor Alexa 555 detector. The interaction was confirmed if we got signal in the FRET channel after subtracting the spectral bleed through. The FRET analysis was done in cellSens Dimension software linked with microscope according to the FRET analysis algorithm by Youvan, et.al. The obtained image after FRET analysis was given in the ratio-metric image panel.

### Fluorescence Assisted Cell Sorting (FACS)

The cells were collected from the plate/flask by trypsinization. After washing once with PBS, they were filtered through .45um nylon membrane. While running through FACS machine, proper controls were kept for setting the gate. For sorting, collection tubes filled with 20% DMEM+ anti-biotic-mycotic cocktail were placed in the collection port. After the sorting was finished, the tubes were centrifuged at 2000 rpm for 10 minutes at room temperature. The cell pellet was further processed based on the experiment.

### H&E Staining

The cryosections were brought to room temperature and dipped in distilled water for 10 mins. It was further semi-dried and dipped in hematoxylin dye for 5 minutes after which the blueing was done in tap water for 10 mins with constant changing of tap water until the whole excess hematoxylin is gone. Then it was dipped in eosin dye for 1 min. Then the slide was dipped in 90% and 100% isopropanol till the red colouring is optimum. It was further fixed with xylene two changes each in an interval of 30 mins. The stained sections were further mounted with DPX and coverslip and observed under light microscope.

### *In silico* analysis of oral cancer patient data

To analyze the pattern of expression of TRIM33 mRNA in oral cancer specimens, data was downloaded from the Gene Expression Omnibus data portal (https://www.ncbi.nlm.nih.gov/geo/). The GEO accession numbers of the datasets are GSE136037, GSE65858. Platform used for expression profiling in all the three datasets were microarray. In GSE136037, histologically confirmed primary tumor of 49 HNSCC patients were divided into 24 without metastasis (“T-without”) and 25 with distant metastasis (“T-with”). In GSE65858 a total of 300 samples were considered for analysis and quality control procedures were applied to microarray probe-level intensity files. A total of 270 tumor arrays remained after removing low-quality arrays, duplicate arrays, and arrays from non-HNSCC samples. Expression values were log2-transformed and normalized using RSN. The relative normalized log transformed microarray data was used for all the analysis. The TCGA data (N=490) was accessed from the TCGA Research Network: https://www.cancer.gov/tcga (accessed on 15 November 2021).

Descriptive statistics were used for all clinical variables. The difference in gene expression levels was evaluated by the Mann–Whitney U test/Kruskal–Wallis test. Correlations were evaluated by Pearsons’s rank test. Heatmap analysis was also performed to analyze association of various genes (TRIM33, TRRAP, HES1, MYC1) with disease free status of the patients. Kaplan-Meier analysis was used to examine the estimated differences in disease-free survival and overall survival between the various groups. Log-rank test (Mantel-Cox) was used to compare the survival between groups. Survival analysis was performed by stratifying tumors based on the upper or third quartile expression of TRIMM33 mRNA levels and grouped into tumors with high and low expression. For all tests, a p-value of <0.05 was considered to be statistically significant. All statistical analysis was carried out using the software XLSTAT - version 2022.2.1.

### Tissue microarray and DAB staining

Samples were collected from Cochin Cancer Research Centre (CCRC). Patients were categorized as recurrent and disease-free groups where patients with disease-free survival of less than 15 months were considered as recurrent (25 samples) while patients with disease-free survival for more than 15 months were considered as disease-free group (27 samples). Immunohistochemistry for the antibodies was done on each of the TMA sections as per standard protocol using the Ventana BenchmarkXT staining system (Ventana Medical Systems, Tucson, AZ, USA). Briefly, 5 μm thick sections were fixed in a hot air oven at 60°C for 60 min and loaded onto the IHC staining machine. De-paraffinization was performed using EZ Prep solution (Proprietary-Ventana reagent), and antigen retrieval was done using Cell Conditioning solution 1 (CC1) for 60 min. The primary antibody was added manually and incubated for 32 min at room temperature. Optiview DAB Detection Kit (Ventana Medical Systems) was used to visualize the signal, using DAB (3–3′diaminobenzidine) as the chromogen. Further, the sections were automatically counterstained with hematoxylin II (Ventana Medical Systems) for 12 min. The slides were removed from the autostainer, washed in de-ionized water, dehydrated in graded ethanol, cleared in xylene, and examined by microscopy. Appropriate positive and negative controls were run for each batch. Two pathologists scored the staining independently. Nuclear/cytoplasmic staining in more >1% of tumor cells was considered positive. Data from 2 recurrent and 26 disease-free samples were used for the statistical analysis.

### RNA-seq Gene Expression Profiling and Pathway Analysis

TIF1γ knockdown and control HSC-4 cells were cultured in sphere medium and RNA was isolated using RNeasy kit (Qiagen). RNA-seq profiles of triplicate samples for each condition were generated from Illumina Nova Seq 6000. Raw RNA-sequencing reads were first assessed for quality using FastQC1 v0.11.8, followed by adaptor and low-quality read removal using fastp2 v1.0.1. Overall quality metrics were summarized with MultiQC3 v1.26. Cleaned reads were aligned to the human reference genome GRCh38 using STAR4 v2.7.3a. Library strandedness was evaluated using RSeQC5 v5.0.4. Gene-level read quantification was performed using FeatureCounts6 v2.0.3 with appropriate strandedness settings. Differential expression analysis was carried out using DESeq27 v1.50.2, applying standard normalization and shrinkage procedures. Gene set enrichment analysis (GSEA)8 was performed using the fgsea package with gene sets obtained from MSigDB. All visualizations, including heatmaps, volcano plots, and enrichment plots, were generated using ggplot2, enrichplot, EnhancedVolcano, and pheatmap.

### Statistical Analysis

All the graphical representations were drawn using GraphPad Prism 8.4. The p-value was calculated by either two-tailed unpaired students t-test or multiple t-test for more than two variables in the GraphPad. * Means p-value<0.05, ** means p-value<0.01, *** means p-value<0.001 and **** means p-value<0.0001 in all the graphs. Kaplan-Meier analysis was also done using the GraphPad and p-value was calculated by log-rank test.

